# Chondroitin 4-sulphate depletion enhances synaptic plasticity and memory in aging

**DOI:** 10.64898/2026.01.28.702127

**Authors:** Jana Svobodova Burianova, Jan Svoboda, Jiri Ruzicka, Katerina Stepankova, Tereza Klausova, Lenka Gmiterkova, Tereza Spundova, Noelia Martinez-Varea, Michaela Kralikova, Rostislav Turecek, Lucia Machova Urdzikova, James W Fawcett, Pavla Jendelova, Jessica CF Kwok

## Abstract

Perineuronal nets (PNNs), specialised extracellular matrix structures enriched in chondroitin sulphate proteoglycans (CSPGs), are key regulators of synaptic plasticity, learning, and memory. Aging is characterised by a shift in chondroitin sulphate composition toward increased chondroitin-4-sulfation (C4S) and reduced C6S, a pattern associated with declining cognitive flexibility. Here, we investigated how selective reduction of C4S affects PNN structure, PV-interneuron connectivity, and cognitive performance across the lifespan. Conditional deletion of the C4-sulfotransferase *Chst11* markedly reduced C4S levels and diminished dendritic PNN complexity while preserving somatic PNN structure. This partial destabilisation of PNNs increased excitatory synaptic input onto PV interneurons in both young and aged mice, without major alterations in basal hippocampal transmission or long-term potentiation. Behaviourally, *Chst11* knockout mice showed robust and persistent protection against age-related cognitive decline. Working memory performance remained stable across aging, short-term spatial memory was enhanced from early adulthood onward, and object recognition memory was significantly prolonged at all retention delays, even in old age. Sociability and social novelty preference were also preserved longer in aging knockouts compared with controls. These improvements occurred despite an overall preservation of PNN architecture, indicating that modifying sulphation rather than removing CSPGs is sufficient to enhance plasticity. Our findings demonstrate that reducing C4S through *Chst11* deletion confers long-lasting enhancements in cognitive function and mitigates aging-related decline. Targeting CS-GAG sulphation patterns may therefore represent a promising strategy for maintaining cognitive resilience and restoring plasticity in aging or neurodegenerative conditions.

## Introduction

The brain extracellular matrix (ECM) is a dynamic structure surrounding neurons and glial cells and is increasingly recognised as a key regulator of brain plasticity throughout life and in neuropathology. Perineuronal nets (PNNs) have attracted particular attention, as specialised ECM structures enriched in chondroitin sulphate proteoglycans (CSPGs, lecticans) that encapsulate specific neuronal populations, predominantly fast-spiking parvalbumin-positive interneurons (Kwok, 2015). Removal of PNNs has been shown to re-activate juvenile-like plasticity and enhance neuroplasticity in the adult brain (Lensjø et al., 2017; Romberg et al., 2013).

PNNs are formed at the end of the critical period during brain maturation (Hou et al., 2017), restricting synaptic remodelling for learning and memory (Beurdeley et al., 2012). Their hierarchical assembly involves hyaluronan, hyaluronan and proteoglycan link proteins (HAPLNs), tenascins, and CSPGs. While PNN formation limits plasticity, ageing and pathological conditions such as neurodegeneration or injury to the central nervous system induce further changes in the neural ECM (Beurdeley et al., 2012; Bernard & Prochiantz, 2016; Testa et al., 2019).

A key determinant of PNN function is the sulphation pattern of chondroitin sulphates (CSs). We have previously shown that chondroitin 4-sulphates (CS-A, primarily via sulfotransferase Chst11) are the predominant form of CS in mature and aged brains and have been shown to be inhibitory to neurite extension and synapse formation (Foscarin et al, 2017). On the other hand, chondroitin 6-sulphates (CS-C, primarily via sulfotransferase Chst3) are expressed at high levels during development and downregulated upon maturation (Foscarin et al, 2017). Over-expression of C6S is able to reactivate plasticity and enhances memory (Yang et al, 2021). Enzymatic digestion of CSPGs using chondroitinase ABC (ChABC) transiently reinstates juvenile-like plasticity, improving memory in aged mice and Alzheimer’s models (Romberg et al., 2013; Yang et al., 2015). Similarly, knockdown of a CSPG aggrecan enhances recovery after synaptic loss (Rowlands et al., 2018; Ruzicka et al., 2022).

Emerging evidence highlights C4S as a critical inhibitory factor. Removal of C4S enhances axonal regeneration and functional recovery after CNS injury (Cheung et al., 2017; Sahu et al., 2019). Furthermore, C4S-rich PNNs in hippocampal CA2 restrict social memory; reducing C4S in this region promotes synaptic plasticity and sociability (Huang et al, 2023). These findings suggest that C4S is not only a barrier to structural remodelling but also a modulator of complex behaviours.

In this study, we hypothesised that sustained C4S abundance in aged brains contributes to cognitive inflexibility for learning and memory, and that reducing C4S can rescue the deficits. Using a developmental *Chst11* knockout model, we target the principal enzyme responsible for C4S synthesis. In these mice with reduced C4S sulphation, we first assessed synaptic connectivity in PNN-associated parvalbumin interneurons and hippocampal electrophysiology. We then evaluated behavioural consequences in learning, memory, and sociability across lifespan stages. We found that reducing C4S through *Chst11* deletion produced a selective loosening of PNN architecture, characterised by reduced dendritic complexity and diminished aggrecan incorporation while preserving somatic PNN organisation. This structural modulation was accompanied by increased excitatory synaptic input onto PV interneurons and age-stable levels of hippocampal synaptic function. Behaviourally, *Chst11* knockout mice exhibited a broad and remarkably enduring enhancement of cognitive performance, including preserved working memory, improved short-term spatial memory, and prolonged object recognition across retention delays, with several of these benefits persisting into late ageing. Social interaction and social novelty discrimination were likewise maintained for longer in *Chst11* knockouts than in controls. Together, these findings demonstrate that targeted reduction of C4S—not wholesale PNN removal—is sufficient to promote synaptic remodelling and sustain cognitive flexibility throughout life, highlighting sulphation-specific modulation of the ECM as a promising strategy for mitigating age-related cognitive decline.

## Material and Methods

### Animals

For all experiments, *Chst11-nestin* knockout (*Chst11*-nestin ko) and *Chst11*-*parvalbumin* knockout (*Chst11*-PV ko) mice were used. The knockout was generated by crossbreeding *Chst11 floxP* mice with either a *nestin-Cre* driver line (Nestin-CreErt2 x (Gt(Rosa)26Sor(CAG-Flpo,-EYFP) x C57BL/6N-A<tm1Brd> Chst11<tm1a(KOMP)Wtsi>/WtsiFlmg)) or *parvalbumin-Cre* driver mice (B6.129P2-Pvalb<tm1(cre)Arbr>/J x (Gt(Rosa)26Sor(CAG-Flpo,-EYFP) x C57BL/6N-A<tm1Brd> Chst11<tm1a(KOMP)Wtsi>/WtsiFlmg)), resulting in either CNS-specific deletion of *Chst11* or deletion of *Chst11* in parvalbumin neurons only. All mice were derived from a C57BL/6 background. Both male and female animals were used throughout the study.

For the behavioural assessment of young animals (6 months; Group 1), four genotypes were included: *Chst11*-*nestin* ko, *Chst11*-*PV* ko, *floxP* littermate controls, and non-littermate wild-type (WT) controls. The same genotype groups were used in the aging (12 months; Group 2) and aged (20 months; Group 3) cohort for behavioural testing.

For biochemical analyses, the left-brain hemisphere was collected from *Chst11*-nestin ko, *Chst11-PV* ko, floxP, and WT mice. The same genotypes were used for immunohistochemical studies.

For electrophysiological recordings of field excitatory postsynaptic potentials (fEPSPs) in acute hippocampal slices, 3–4 animals per genotype (*Chst11-nestin* ko, *Chst11-PV* ko, *FloxP*, WT) were examined at 3 and 22 months of age.

All experiments were performed in accordance with the European Communities Council Directive of 22 September 2010 (2010/63/EU), regarding the use of animals in research and were approved by the Ethics Committee of the Institute of Experimental Medicine ASCR, Prague, Czech Republic.

### Tissue isolation and preparation

For immunohistochemistry, the mice were transcardially perfused with PBS followed by paraformaldehyde (4% PFA in PBS). The brains were left in PFA overnight then gently washed out in PBS, 20% sucrose and then 30% sucrose (each solution time= 24 h, t=4°C). Using cryotome, 20 µm thick coronal sections were prepared.

For GAG biochemistry, the whole brain tissue was isolated on the dry ice and then stored at -80°C until further use.

### Immunohistochemistry

To observe integrity and morphology of PNNs in *Chst11* ko, staining biotinylated *Wisteria floribunda* agglutinin (WFA, 1:150, Sigma-Aldrich, Germany), and anti-aggrecan primary antibody (1:150, Sigma-Aldrich) together with DAPI (1:1000, Invitrogen) were used. For analysis of parvalbumin positive inhibitory interneurons synaptic connectivity, antibodies against parvalbumin (1:50, Synaptic Systems), PSD95 (1:100, excitatory postsynaptic, Synaptic Systems), gephyrin (1:100, inhibitory postsynaptic, Synaptic Systems), vGLUT1 (1:100, excitatory presynaptic, Synaptic Systems), and vGAT (1:100, inhibitory presynaptic, Synaptic Systems) were used. To visualize primary antibodies positivity anti-guinea pig Alexa Fluor 594 (1:200, Invitrogen), anti-rabbit Alexa Fluor 594 (1:200, Invitrogen), anti-chicken Alexa Fluor 594 (1:200, Invitrogen) and anti-mouse Alexa Fluor 488 (1:200, Invitrogen) or anti-rabbit Alexa Fluor 405 (1:100, Invitrogen) secondary antibodies were used. To visualize the primary antibody a streptavidin Alexa Fluor 488 (1:300, Invitrogen).

### Confocal imaging and super resolution microscopy

A Zeiss 880 confocal microscope with a 20x objective lens and Z-stack imaging focused on the CA1 region of the hippocampus was used to describe the altered morphology of PNN. A low-magnification objective was used to capture the entire section of neurons covered by PNN. Lasers of 488 nm (WFA), 594 nm (Aggrecan), and 405 nm (DAPI) were used. Laser power was maintained below 3%. (Figure 1, 1S, 2S).

**Figure 1.**
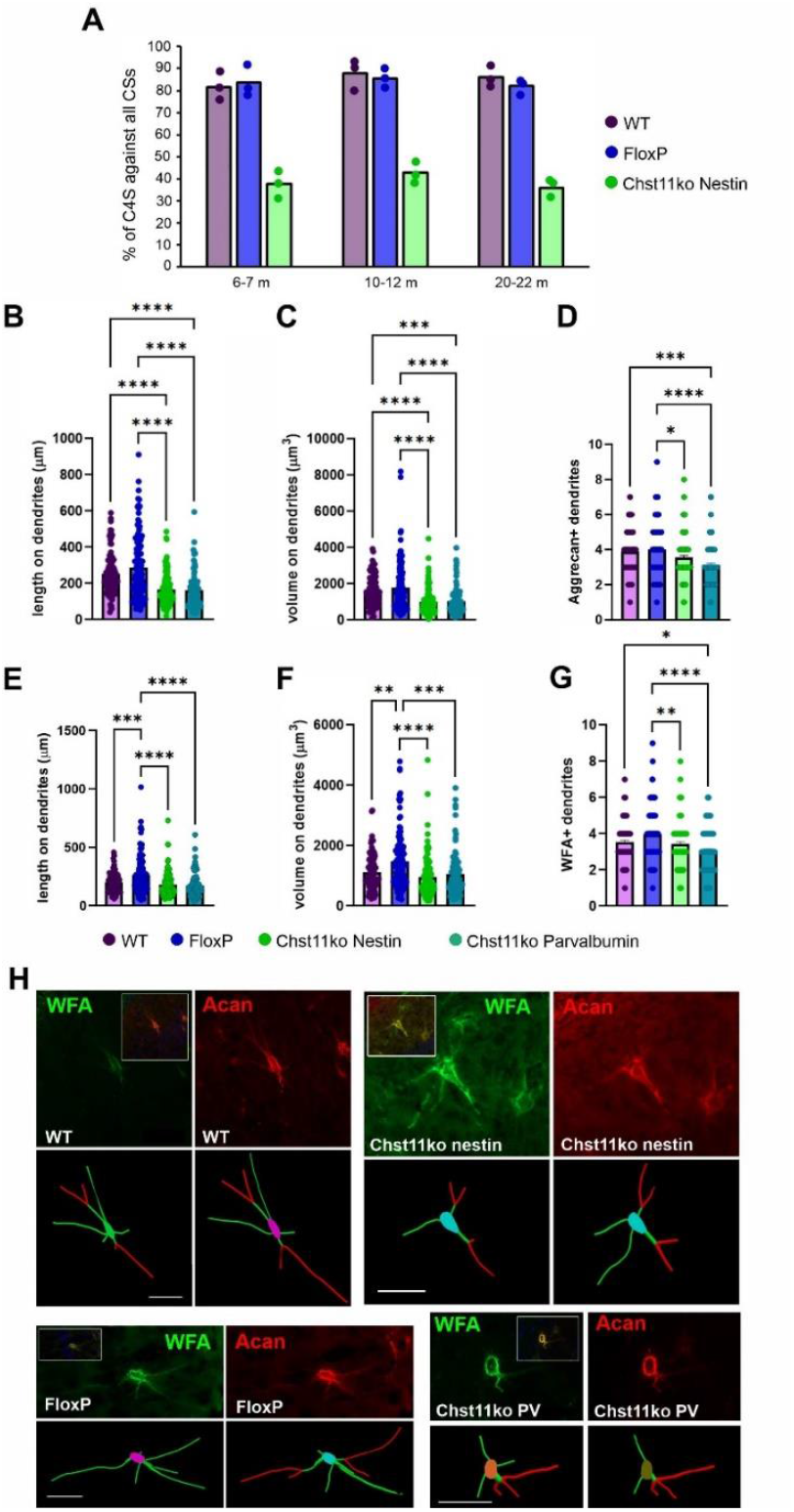
Chst11 knockout reduces chondroitin-4-sulfation and decreases PNN structural complexity and aggrecan incorporation in the hippocampal CA1 region. **(A)** Quantification of chondroitin sulphate sulphation composition in whole-brain glycosaminoglycan (GAG) extracts from WT/FloxP controls and Chst11 knockout mice at young adult, adult, and aged time points. Chst11 knockout results in a robust and sustained reduction of chondroitin-4-sulfation (C4S) across the lifespan. Perineuronal net (PNN) morphology in the CA1 area of the hippocampus was analysed using immunohistochemical labelling for aggrecan (B–D) and Wisteria floribunda agglutinin (WFA; E–G), followed by three-dimensional reconstruction and quantitative analysis using Neurolucida software. Quantified parameters included PNN length on dendrites (B, E), PNN volume on dendrites (C, F), and the number of aggrecan-positive or WFA-positive dendrites (D, G). Data are shown as individual dendritic measurements with bars representing mean ± SEM. Statistical analysis was performed using one-way ANOVA followed by Tukey’s post hoc tests. Statistical significance is indicated as * *p* < 0.05, ** p < 0.01, *** *p* < 0.001, **** p < 0.0001. (G) Representative confocal images and corresponding three-dimensional reconstructions of PNNs labelled with WFA (green) and aggrecan (red) in WT, FloxP, Chst11-nestin knockout, and Chst11-PV knockout mice. Scale bar, 50 μm.

To estimate the connectivity of synapses to parvalbumin-positive (PV^+^) inhibitory interneurons in the CA1 region of the hippocampus, we used a Zeiss LSM 880 confocal microscope with Airyscan detection in super-resolution mode (Figure 2S). Synaptic colocalisation was acquired using a 100× oil-immersion objective (α Plan-Apochromat 100×/1.46 Oil DIC M27 Elyra), and neuronal regions of interest were selected using the zoom function (2x). Airyscan super-resolution imaging was performed using all 32 detectors at the highest SR setting, with three independent channels recorded. Laser power was kept below 15% to preserve fluorophore stability. Synapse quantification was based on the colocalisation of the following markers: presynaptic (vGlut1 or vGAT, 594 nm); postsynaptic (PSD95 or Gephyrin, 488 nm); PV^+^ (405 nm). For each animal, 11–15 PV^+^ neurons were imaged with at least 20 optical slices per neuron (n = 3-4 animals per group).

**Figure 2.**
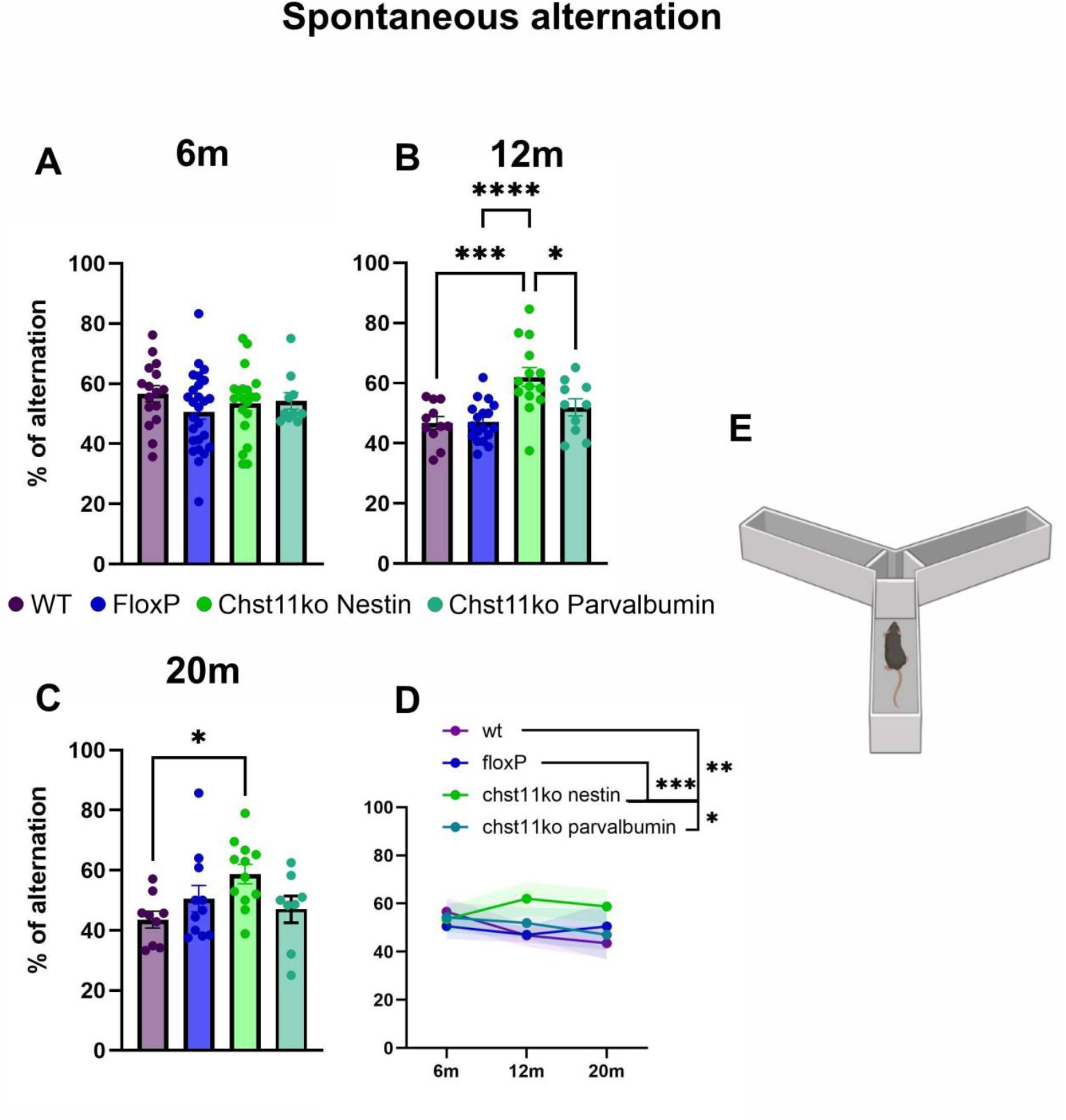
Age-dependent decline in spatial working memory is prevented in *Chst11* ko mice. Spontaneous alternation was assessed using a Y-maze at 6 (A), 12 (B), and 20 (C) months of age. Spatial working memory performance was quantified as the percentage of alternations, defined as consecutive entries into all three arms without repetition. Individual data points represent single animals, with bars indicating mean ± SEM. (D) Line plot summarising changes in spontaneous alternation across ages for each experimental group. Shaded areas represent CI. (E) Schematic illustration of the Y-maze apparatus. * p < 0.05, ** p < 0.01, *** p < 0.001, **** p < 0.0001

Synaptic puncta were quantified using Fiji (ImageJ) by applying a threshold to each channel and utilising the ‘Analyze Particles’ tool to detect presynaptic, postsynaptic and colocalised puncta within PV^+^ cell boundaries. Synapse density was calculated by first estimating the PV^+^ cell volume from the 2D cell area (µm^2^), the number of Z-stack optical planes and the Z-step height (µm). Cell thickness was defined as the number of Z steps multiplied by the step height. Total cell volume (µm^3^) was obtained by multiplying thickness by cell area. The final synapse density (synapses/µm^3^) was then computed by dividing the number of colocalised puncta by the calculated cell volume.

### PNN morphology

Tracing of perineuronal nets was performed using bright-field and fluorescent microscopy (Leica DMRXA microscope) coupled to a high-resolution monitor (Wacom, 3840 ×2160 px) with Neurolucida 2021.1.3 software (MicroBrightField Bioscience, Williston, Vermont, United States). The 3D tracing was performed manually, starting with neuronal soma. Perineuronal nets on dendritic branches were traced with a simultaneous slow movement along the z-axis to keep the traced segment at maximum focus. Twenty neurons per animal (six animals per group) were selected and traced for each staining. The final PNNs tracings of particular neurons were saved and Neurolucida Explorer 2021.1 (MBF Bioscience) was used to conduct a detailed morphological structure analysis. From 3D reconstruction PNNs, the following morphometric parameters were quantified: area of PNN on cell body, length on dendrites, volume of PNN on dendrites, number of dendrites with PNN, number of nodes in PNN sheath, and number of ends. The obtained data were further exported and statistically processed with GraphPad Prism software (9.0). Differences between groups were analysed using one-way ANOVA, followed by Tukey’s multiple comparisons test.

### GAG isolation and quantification

Frozen tissues were weighed before incubation with acetone to remove the lipids. The samples were then homogenised and treated with pronase (15 mg pronase per hemisphere, in 0.1 M Trizma hydrochloride, 10 mM calcium acetate), pH 7.8. Proteins were then removed by precipitation and the supernatant, which contains the GAGs, was collected and stored on ice. The solution was then neutralised with 1 M Na_2_CO_3_ to pH 7.0, and the GAGs were recovered by ethanol precipitation. The isolated GAGs were redissolved in 0.3 ml of deionised water. The GAG concentration was quantified using cetylpyridium chloride (CPC) turbidimetry against a standard curve prepared from chondroitin sulphate A. Briefly, the diluted sample was mixed with 0.2% (w/v) CPC and with 133 mM MgCl_2_ in ratio 1:1. Absorbance was measured at 405 nm using plate reader spectrophotometer (FLUOstar® Omega, BMG LABTECH). Each sample was carried out in three different dilutions and each dilution was carried out in duplicate.

### Acute slices and electrophysiology

Mouse brains were dissected, and 300 um thick parasagittal hippocampal slices were prepared in ice cold solution composed of (in mM): NaCl 130, KCl 3.5, MgCl_2_ 3, CaCl_2_ 0.5, glucose 10, NaH_2_PO_4_ 1.25, NaHCO_3_ 24; supplements were added to the solution oxygenated with 5% CO_2_ at final concentration (in mM): Na pyruvate 4.9, L-ascorbic acid 0.5. After incubation for 45min at 35°C fEPSPs were induced by minimal stimulation of CA3 and recorded in the *stratum radiatum* of CA1. Sixty baseline responses were recorded at stimulation frequency 0.05 Hz before LTP induction by 4 trains delivered at 15s intervals. Each train consisted of 10x 4-pulse bursts (inter-pulse interval of 10ms and inter-burst interval of 200ms). One hundred twenty fEPSP responses were recorded 3 min after the LTP protocol. Linear regression of responses before LTP was used to estimate baseline fEPSP amplitude at the time of 115th LTP response. Amplitude of LTP response was determined as average of the last 10 responses after LTP induction. LTP induced increase in fEPSP amplitude was expressed as percentage of the baseline. fEPSPs were recorded using Axopatch 200B, Axon Digidata 1550B and Clampex 10.7. Analysis of response amplitudes was done in Clampfit 10.7 and GraphPad Prism 10.2 was used for statistical analysis.

### Behavioral experiments

If not stated otherwise, the sequence of the behavioural tasks was as follows: Three-chamber task, Spontaneous alternation, Novel object recognition, Morris water maze.

### Three chamber task - Sociability and social memory

The three-chamber task (Ugo Basile, Italy) was used to assess the impact of Chst11 knockout on sociability and social recognition memory. Animals aged 6, 12, and 20 months were placed in the middle chamber with the sliding doors closed and allowed to explore for 5 minutes. After this habituation period, the doors were opened, and the mice were allowed to freely explore all three chambers.

During the first 10-minute period (sociability phase), one side chamber contained an empty wire cage, while the other housed a “stranger” mouse of the same sex. In the following 10-minute period (social recognition memory phase), a second stranger mouse was introduced into the previously empty cage.

To assess sociability and recognition memory, the ratio of time spent interacting with the occupied versus empty cage, and with the novel versus familiar mouse, respectively, was analysed. The protocol was adapted from Yang et al. (2011).

### Spontaneous alternation test

The spontaneous alternation test in a Y-maze was conducted prior to the novel object recognition task, serving as an adaptation phase to the maze environment. Mice were allowed to freely explore the three-armed maze for 5 minutes. The maze consisted of three symmetrically arranged arms (length: 15 cm), each with opaque walls (height: 30 cm). The sequence of arm entries was recorded from video footage.

Spontaneous alternation performance was calculated as SA% = (# of spontaneous alternations / [total number of arm entries − 2]) × 100, following the method described by Prieur and Jadavji (2019). Additionally, the overall visit frequency was also assessed.

### Spontaneous novel object recognition test

The spontaneous object recognition (SOR) task was performed as previously described (Romberg et al., 2013; Yang et al., 2015), using the same Y-maze apparatus adapted from the spontaneous alternation test. The maze consisted of three arms (length 15 cm), with one designated as the start arm and the other two used for object presentation. For the SOR task, the object arms were shortened to 10 cm using sliding doors to maximize the time spent in the object exploration area. Mice were previously habituated to the apparatus during the spontaneous alternation test, which was conducted at least 48 hours prior to the SOR task.

During the sample phase, mice were allowed to explore two identical objects, each placed in one of the two object arms, for 5 minutes. In the choice phase, both objects were replaced: one with an identical copy and the other with a novel object. After a designated delay period, mice were reintroduced to the maze for 5 minutes to explore the objects. Different delay intervals were used to assess short-term, and long-term, memory (e.g., 3 hours, 24 hours or 48 hours). A minimum of 48 hours separated different sample/choice interval trials.

Object exploration time was assessed from video recordings by two independent examiners. Exploratory behaviour was defined as direct nasal or head contact with an object. The discrimination ratio was calculated as the difference in time spent exploring the novel and familiar objects, divided by the total object exploration time.

### Morris water maze

A short-term spatial memory test, modified from Ruzicka et al. (2016), was conducted. The experiment lasted for five consecutive days, with four trials per day, all starting from the same initial position.

To engage working memory, both the platform location and the starting point were changed each day. Performance in the first trial of each day was set as the baseline (100%), the memory was indexed as the percentage of reduction in latency to locate the hidden platform in trials 2–4 relative to trial. The average ratio across the five days is illustrated in Figure 3.

**Figure 3.**
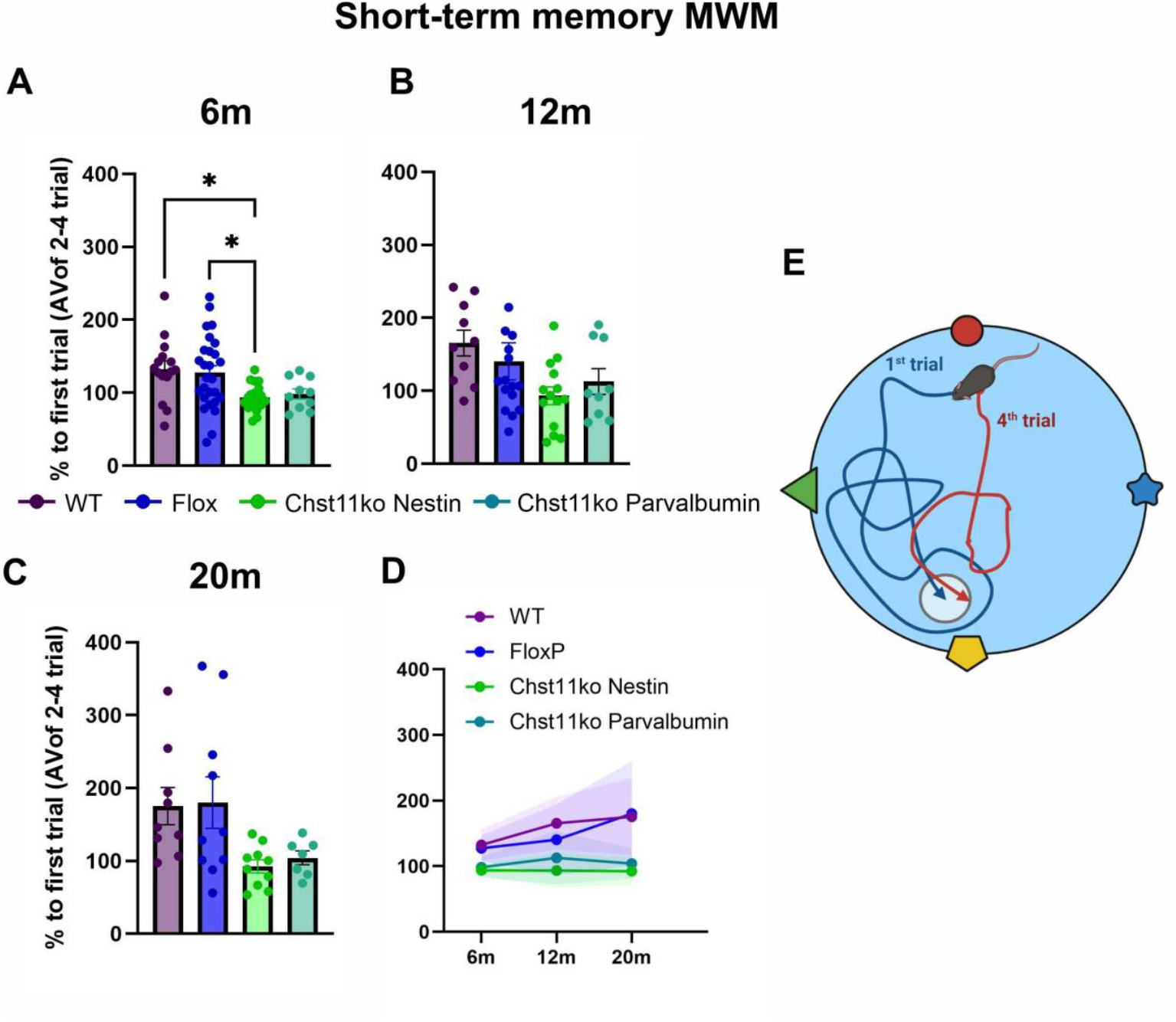
Short-term spatial memory is preserved in Chst11-nestin mice in the Morris Water Maze (MWM). Short-term memory was assessed using a four-trial daily Morris water maze (MWM) protocol at 6 (A), 12 (B), and 20 (C) months of age. Memory performance was quantified as the percentage reduction in latency to locate the hidden platform in trials 2–4 relative to trial 1. Individual data points represent single animals, with bars indicating mean ± SEM. (D) Line plot summarising changes in short-term memory performance across ages for each experimental group, with shaded areas representing CI. (E) Schematic illustration of the Morris water maze task. * p < 0.05, ** p < 0.01, *** p < 0.001, **** p < 0.0001

### Statistics

Statistical analyses were performed using GraphPad Prism (10.6.1) Data distribution was assessed using Shapiro–Wilk tests, and homogeneity of variance was evaluated using Brown–Forsythe and Bartlett’s tests where appropriate. Group differences were analysed using one-way or two-way analysis of variance (ANOVA), depending on the experimental design. For experiments involving two independent factors, two-way ANOVA was applied, followed by post hoc comparisons using Tukey’s or Šídák’s multiple-comparison tests as appropriate. Statistical significance is reported as *p* values, and effect sizes are reported as partial eta squared (ηp^2^) for all ANOVA-based analyses. Post hoc results are described where relevant to indicate the direction of significant effects. Data are presented as mean ± SEM or CI unless stated otherwise. Statistical significance was set at *p* < 0.05.

## Results

### Knockout of *Chst11* leads to reduction in PNN structural complexity and decreases aggrecan incorporation

We first verified the extent of C4S reduction *in Chst11* knockout mice. GAGs were isolated from the brains at three age stages, young adult (6-7 months), adult (10-12 months), and aged (20-22 months old), and the sulphation composition of CSs was quantified (Fig. 1A). C4S levels were significantly reduced from over 80% in WT and *FloxP* controls, to 38% in young adults, 43% in adults, and 36% in aged Chst11 knockout animals.

To assess the impact of reduced CSPG sulphation on PNN morphology at the conclusion of the first behavioural phase (7 months of age), we performed three-dimensional reconstruction and quantitative analysis of Z-stack confocal images using Neurolucida software on CA1 PNN-positive neurons. Key morphometric parameters included PNN length on dendrites, PNN volume on dendrites, and the number of dendrites covered by PNNs. Two complementary markers were used for visualisation, *Wisteria floribunda* agglutinin (WFA) and anti-aggrecan antibody, which share overlapping yet distinct binding profiles (Miyata & Kitagawa, 2016; Irvine & Kwok, 2018; Rowlands et al., 2018). Analysis revealed a pronounced reduction in PNN complexity in *Chst11* knockout mice compared to *FloxP* littermate controls and wild-type animals (Fig. 1B-H). Specifically, quantitative three-dimensional analysis demonstrated a significant genotype-dependent reduction in PNN length on dendrites, PNN volume on dendrites, and the number of PNN-positive dendrites. Using aggrecan immunolabelling, genotype exerted a strong effect on PNN length on dendrites (*p* < 0.0001, ηp^2^ = 0.16) and PNN volume on dendrites (*p* < 0.0001, ηp^2^ = 0.12), with a more moderate but still significant effect on the number of aggrecan-positive dendrites (*p* < 0.0001,ηp^2^ = 0.06).

Consistent results were obtained using Wisteria floribunda agglutinin (WFA) staining, which also revealed significant genotype effects on PNN length on dendrites (*p* < 0.0001, ηp^2^ = 0.09), PNN volume on dendrites (*p* < 0.0001, ηp^2^ = 0.07), and the number of WFA-positive dendrites (*p* < 0.0001, ηp^2^ = 0.06).

Across both markers, the largest effects of Chst11 deletion were observed for PNN length and volume on dendrites, whereas reductions in the number of PNN-positive dendrites were comparatively smaller, indicating a preferential destabilisation of dendritic PNN continuity rather than a complete loss of dendritic PNN coverage.

### Working memory is preserved in early aging by transgenic reduction of C4 epitope sulfation

To determine the impact of the C4S sulphation epitope on spatial working memory, we performed the Y-maze spontaneous alternation (SA) test across different ages. At 6 months, all groups including WT, *FloxP* controls, and *Chst11* knockout mice displayed comparable performance, indicating normal behaviour in young animals (Fig. 2A). In aging cohorts, a significant decline in SA performance was observed in both control groups at 12 months (Fig. 2B, group effect *p*<0.0001, ηp^2^=0.36), whereas *Chst11-nestin* knockout mice maintained higher alternation rates. At 20 months, this protective effect persisted, with *Chst11*-*nestin* knockout mice performing significantly better than controls (Fig. 2C, D group effect *p*=0.0375, ηp^2^=0.21). Spontaneous alternation was quantified as the percentage of consecutive entries into all three arms without repetition. Line graphs summarize group trajectories across ages (Fig. 2D). A two-way ANOVA revealed a significant age × genotype interaction (p=0.0138, ηp^2^ = 0.09), driven by the selective improvement in *Chst11*-*nestin* ko mice at 12 and 20 months. No main effect of age was observed (p=0.23), whereas a robust main effect of genotype emerged (p=0.0004, ηp^2^ = 0.10)).

Overall, these findings demonstrate that age-dependent decline in spatial working memory is prevented in *Chst11* knockout mice.

### Mice with decreased C4S epitope, express enhanced and preserved short-term memory

In order to determine the impact *Chst11*ko on spatial short-term memory, the Morris water maze test was used. At the 6 months of age, *Chst11-nestin* ko animals have shown significantly enhanced short term memory performance in MWM in comparison to *FloxP*, WT control groups and *Chst11-PV* ko group (Fig. 3, genotype effect *p*=0.0062; ηp^2^=0.17). As the mice aged, the worsening of performance in short-term memory in MWM was observed. At 12 months, group effect did not reach significance. At 20 months, genotype significantly affected performance (*p*=0.0215; ηp^2^≈0.26) despite heterogeneity of variances (Bartlett *p*=0.0001). However, no post-hoc comparison survived multiple-comparisons correction, likely reflecting increased inter-animal variability. *Chst11*-*nestin* ko and *Chst11-PV* ko both tended to perform better, but effects remained non-significant.

To assess performance across aging, a two-way ANOVA including all animals (factors: group × age) showed no interaction (p=0.602) and no main effect of age (p=0.118), but a robust main effect of genotype (p<0.0001; ηp^2^=0.18) on short-term spatial memory, with consistently superior performance in *Chst11*-*Nestin* ko mice across the lifespan (Fig.3 C, D, E).

### Object based long-term memory is enhanced by C4S reduction

Spontaneous novel object recognition relies on the integrity of the perirhinal cortex and hippocampus. Previous studies have shown that PNN digestion or modification can prolong exploration of previously encountered objects (Romberg et al., 2013; Yang et al., 2015; Yang et al., 2017; Yang et al., 2021). In our study, we examined the impact of PNN modification on the persistence of object memory across three retention delays (3 h, 24 h, 48 h) and three age points (6, 12 and 20 months). Across all ages, object recognition performance in all delays was strongly influenced by genotype, whereas neither delay nor the delay × group interaction reached statistical significance at any age. At 6 months, the effect of group on the discrimination index was relatively strong (p<0.0001, ηp^2^=0.30), reflecting enhanced object memory in both *Chst11-nestin* ko and *Chst11-PV* ko mice compared with WT and especially *FloxP* controls (Fig. 4A). At 12 months, group differences in the discrimination index remained pronounced (p<0.0001, ηp^2^=0.35). Unlike WT and *FloxP* mice—whose performance declined toward chance—both *Chst11*-*nestin* ko and *Chst11*-*PV* ko lines continued to show clearly superior object recognition memory across delays (Fig. 4B). At 20 months, the effect of group on the discrimination index remained significant (p<0.0001, ηp^2^=0.24), driven primarily by persistently high discrimination in *Chst11-nestin* ko mice. In contrast, *Chst11-PV* ko mice no longer outperformed WT or *FloxP* controls and showed discrimination indices close to, or in some delays below, control levels (Fig. 4C). Taken together, these findings demonstrate a remarkably stable enhancement of object recognition memory in *Chst11-nestin* ko mice across the lifespan — a profile that persists even at 20 months with a 48-hour delay, when memory performance in all other groups has declined toward chance.

**Figure 4.**
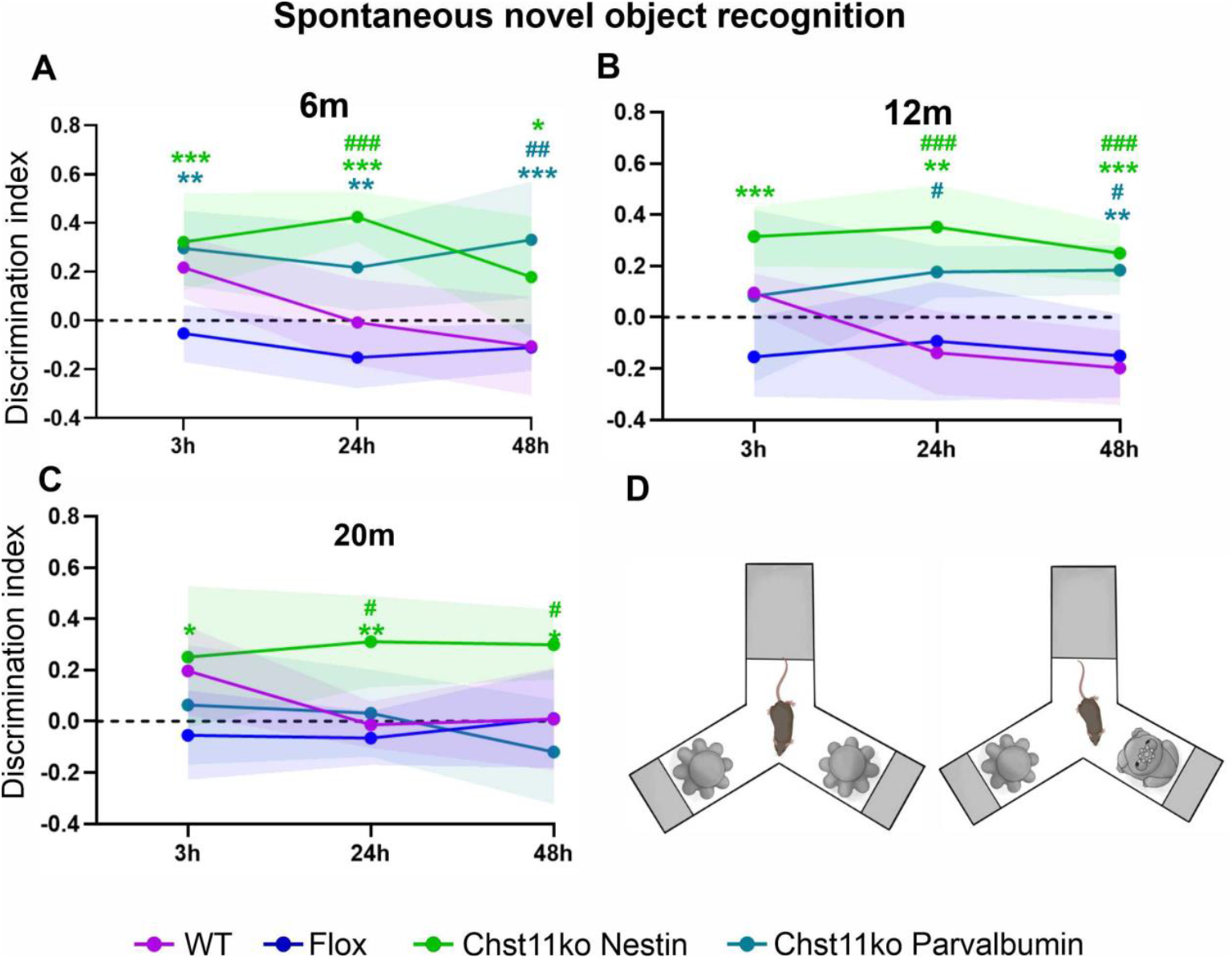
Long-term object recognition memory is preserved in *Chst11*ko Nestin and *Chst11*ko Parvalbumin mice. Spontaneous novel object recognition was assessed using the discrimination index at different delays (3, 24, and 48 h) at 6 (A), 12 (B), and 20 (C) months of age. The dashed line indicates chance performance. Data are shown as mean ± CI. Green asterisks and hashtags indicate statistical significance of the Chst11-nestin knockout group relative to WT and FloxP groups, respectively. Blue asterisks and hashtags indicate statistical significance of the Chst11-PV knockout group relative to WT and FloxP groups, respectively. (D) Schematic illustration of the familiarization (left) and test (right) phases of the novel object recognition task.

### Sociability and social memory are improved and preserved in early ageing in animals with reduced C4 epitope sulfation

The impact of *Chst11* ko on sociability and social novelty was measured using the three-chamber test across 6, 12 and 20 months of age. After habituation phase, the animals were allowed to explore the whole apparatus with chamber containing a cage with a stranger mouse and the second chamber containing an empty cage. At 6 months, only WT and *Chst11-nestin* ko mice showed significant sociability, spending more time interacting with the social stimulus than the empty cage (Fig. 5A, main effect of Cage p=0.0003, ηp^2^=0.18; effect of Group *p*=0.008, ηp^2^=0.14; interaction *p*=0.001, ηp^2^=0.21). At 12 months, significant sociability appeared in both *Chst11* knockout lines, whereas WT and *FloxP* controls showed no preference for the conspecific (Cage p=0.001, ηp^2^=0.20; Group p=0.002, ηp^2^=0.15; Cage × Group interaction p = 0.0011, ηp^2^ = 0.30; Fig. 5B). At 20 months, sociability remained evident primarily in *Chst11-nestin* ko and *Chst11-PV* ko mice, whereas WT and *FloxP* showed weaker or absent social preference (Fig. 5C).

**Figure 5.**
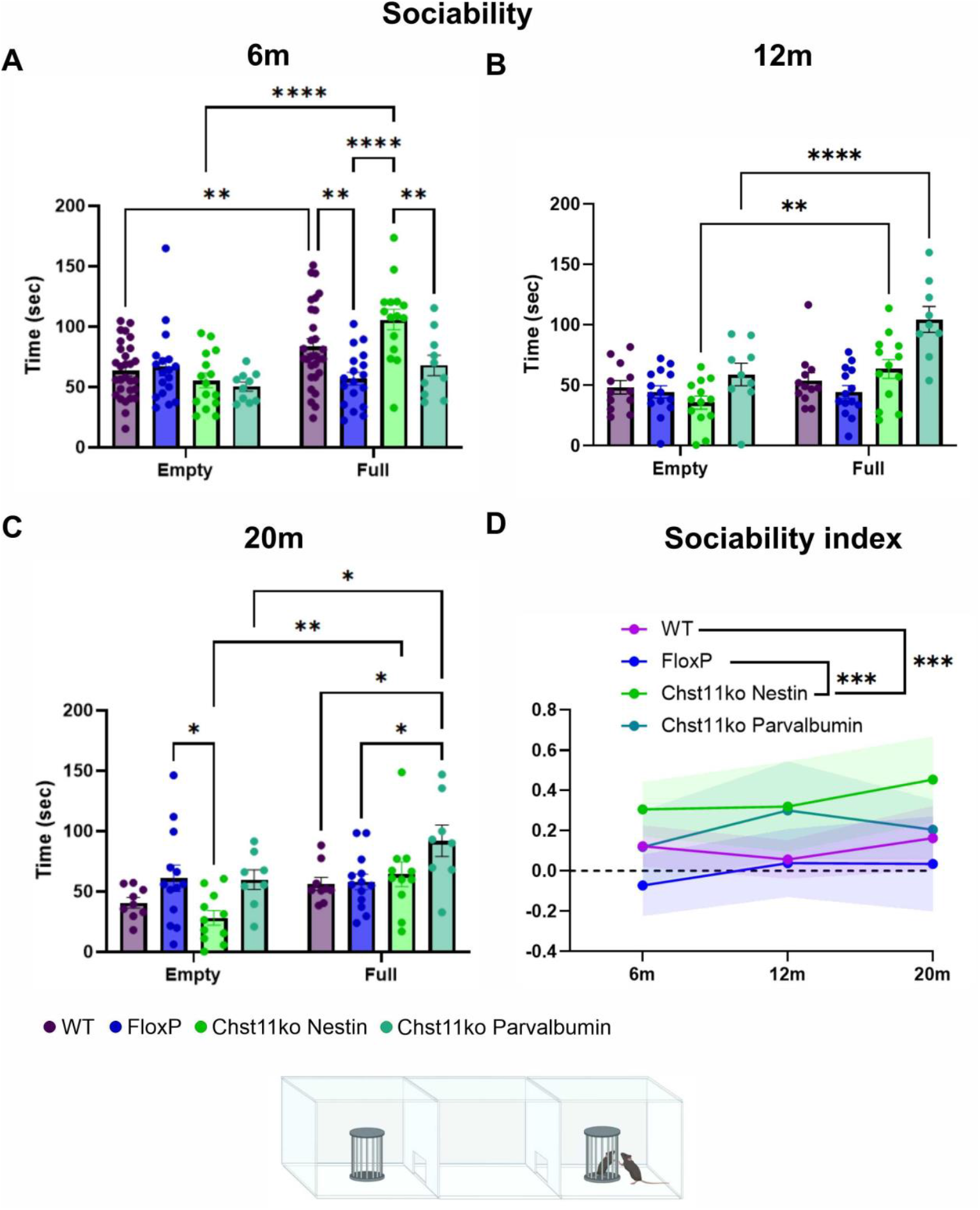
Sociability deficits are rescued in *Chst11-nestin* ko and *Chst11-PV* ko mice. Sociability was assessed using the three-chamber test at 6 (A), 12 (B), and 20 (C) months of age. Mice were given the choice between a chamber containing a stranger mouse (Full) and an empty chamber (Empty), and the time spent in each chamber was recorded. Individual data points represent single animals, with bars indicating mean ± SEM.(D) Sociability index across ages, calculated as the difference between time spent in the Full and Empty chambers relative to total exploration time. Shaded areas represent CI.

Immediately after the sociability test, a second stranger was introduced, allowing assessment of preference for the novel versus familiar mouse. (Fig. 6). At 6 months of age WT, *Chst11-nestin* ko, and *Chst11-PV* ko mice showed robust novelty preference, whereas *FloxP* did not. (Fig. 6A; Novelty main effect p < 0.0001, ηp^2^ = 0.39; Group × Novelty interaction p = 0.0002, ηp^2^ = 0.25). At 12 months, novelty preference persisted in both *Chst11* knockout groups, but not in the control groups (Fig. 6B; significant effect of interaction p = 0.011, ηp^2^ = 0.16. By 20 months, novelty discrimination declined across groups and no longer differed significantly between genotypes (Fig. 6C).

**Figure 6.**
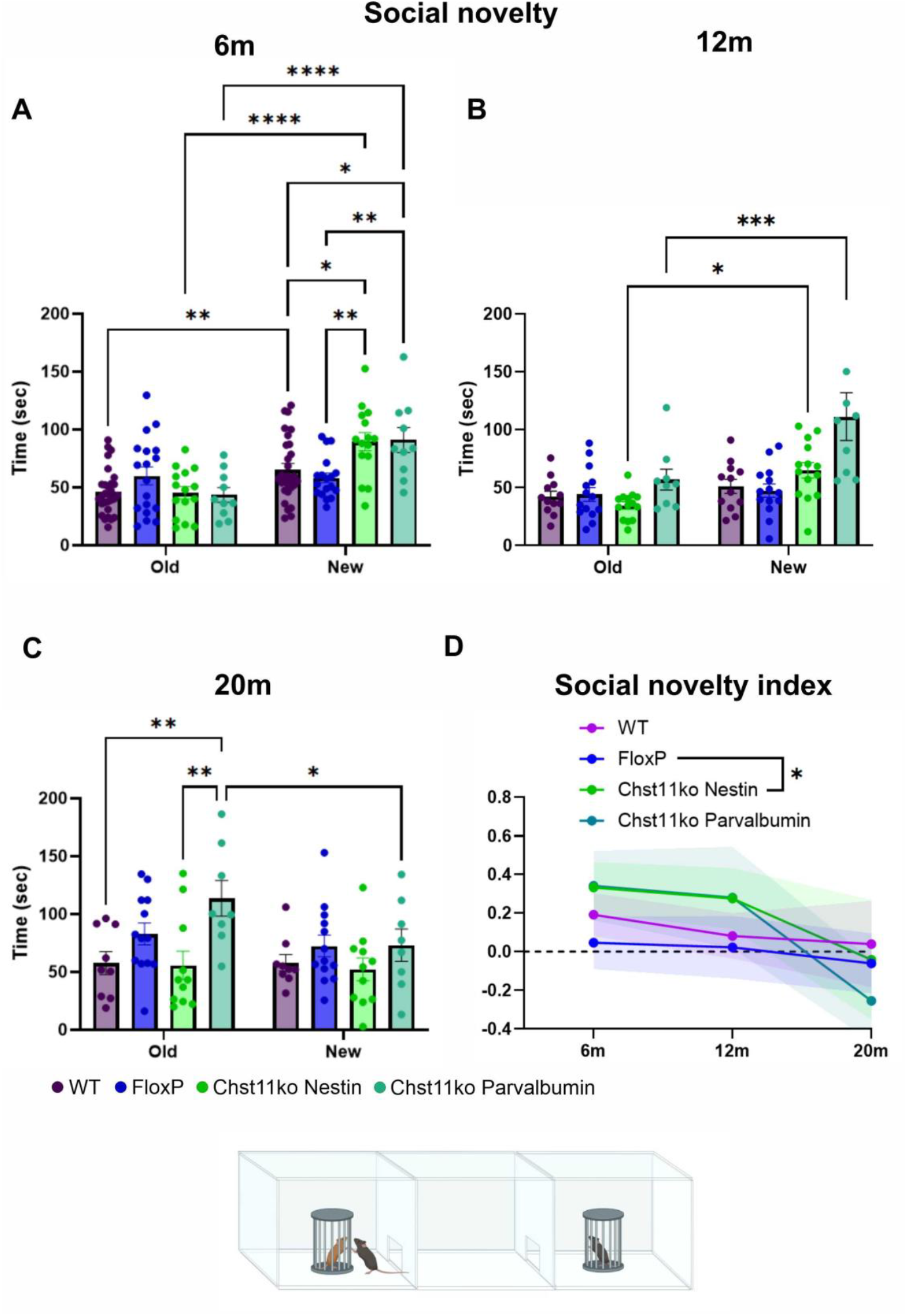
Social novelty preference is preserved in *Chst11-nestin* ko and *Chst11-PV* ko mice. Social novelty was assessed using the three-chamber task at 6 (A), 12 (B), and 20 (C) months of age by measuring the time spent interacting with a novel mouse (New) versus a familiar mouse (Old). Individual data points represent single animals, with bars indicating mean ± SEM. (D) Social novelty index across ages, calculated as the difference between time spent with the novel and familiar mouse relative to total exploration time. Shaded areas represent CI.

Across ages, the sociability index was strongly affected by Group (p<0.0001, ηp^2^=0.20) but not Age. The social novelty index showed significant Age effects (p=0.001, ηp^2^=0.16) and a modest Group effect (p=0.343, ηp^2^=0.05), without a significant interaction. These indices corroborate that *Chst11-*ko mice maintain social motivation and novelty preference longer during aging, whereas control groups show an age-related decline. (Fig. 5D, 6D).

### Reduction of C4S increases synaptic input on PV^+^ interneurons but no change in field EPSPs

Chst11 knockout markedly reduces the structural complexity of PNNs and aggrecan incorporation in hippocampal CA1 neurons. To determine how this reduction in PNN integrity affects synaptic connectivity, we next investigated synaptic inputs onto parvalbumin-positive interneurons in the hippocampus. The co-localisation of pre and postsynaptic markers on the soma of PV neurones was performed using single-neuron Z-stack images acquired with Airyscan super-resolution microscopy. Excitatory synapses were identified by the co-localisation of vGlut1 and PSD95, while inhibitory synapses were identified by the co-localisation of vGAT and gephyrin (Fig. 7A, B).Excitatory synaptic input onto PV^+^ interneurons differed significantly between genotypes at both 6 and 20 months of age (6M: p < 0.0001, ηp^2^ = 0.25; 20M: p < 0.0001, ηp^2^ = 0.29). Post hoc analyses revealed that both Chst11-PV and Chst11-nestin knockout mice exhibited significantly increased excitatory synapse density compared with WT and FloxP controls (Fig. 7A). Inhibitory synaptic input onto PV^+^ interneurons showed an age-dependent and genotype-specific pattern. At 6 months of age, inhibitory synapse density differed modestly between genotypes (*p* = 0.0147, ηp^2^ = 0.06), with post hoc analysis indicating a small increase in Chst11-PV knockout mice compared with FloxP controls (Fig. 7C). In contrast, at 20 months of age, inhibitory synaptic input differed strongly between genotypes (*p* < 0.0001, ηp^2^ = 0.31). Post hoc analyses revealed a pronounced increase in inhibitory synapse density in Chst11-PV knockout mice, whereas Chst11-nestin knockout mice exhibited a significant reduction compared with both WT and FloxP controls (Fig. 7D).

**Figure 7.**
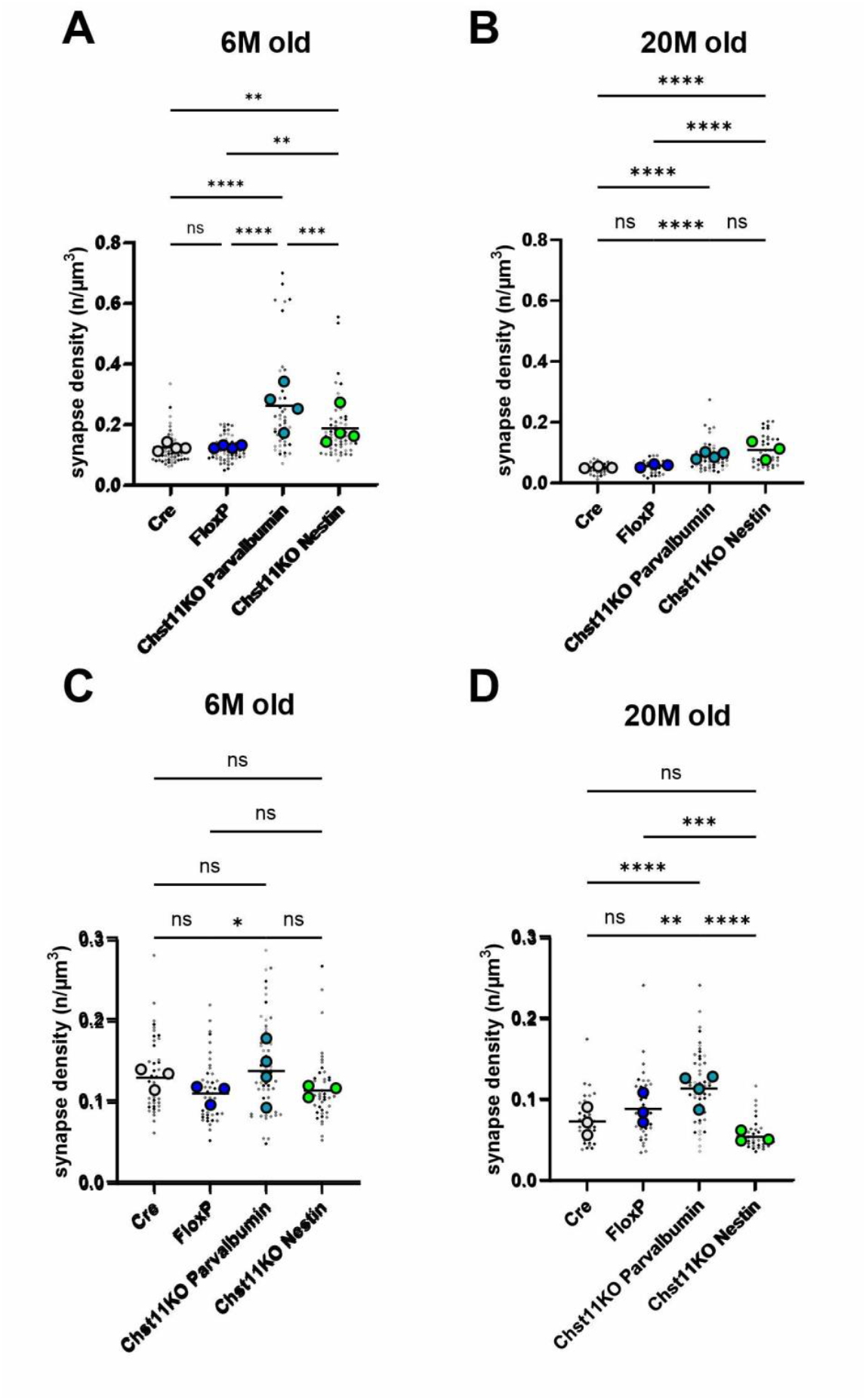
Reduced C4S alters synaptic input onto PV^+^ interneurons in hippocampal CA1. Synaptic input onto parvalbumin-positive (PV^+^) interneurons was quantified from Airyscan z-stacks as colocalised puncta of vGlut1/PSD95 (excitatory) or vGAT/gephyrin (inhibitory) on PV^+^ somata and expressed as synapse density (puncta/µm^3^). (A, B) Excitatory synapse density at 6 months (A) and 20 months (B). (C, D) Inhibitory synapse density at 6 months (C) and 20 months (D). Small dots represent individual PV^+^ neurons; larger coloured symbols indicate animal means. Data are shown with mean. One-way ANOVA with Tukey’s post hoc test. *p < 0.05, **p < 0.01, ***p < 0.001, ****p < 0.0001.

To assess the functional consequences of altered synaptic connectivity, we examined long-term potentiation (LTP) in the CA3–CA1 pathway using acute hippocampal slices. LTP magnitude, quantified as the percentage change in fEPSP amplitude relative to baseline, differed significantly between experimental groups (*p* = 0.0016, ηp^2^ = 0.35). Post hoc analysis revealed a pronounced increase in LTP in young (3-month-old) Chst11-PV knockout mice compared with age-matched WT and Chst11-nestin knockout mice (Fig. 8B). In contrast, significant differences in LTP magnitude were detected between genotypes at 20 months of age, with LTP levels in aged Chst11-PV knockout mice being lower to those observed in age-matched WT (Fig. 8E).

**Figure 8:**
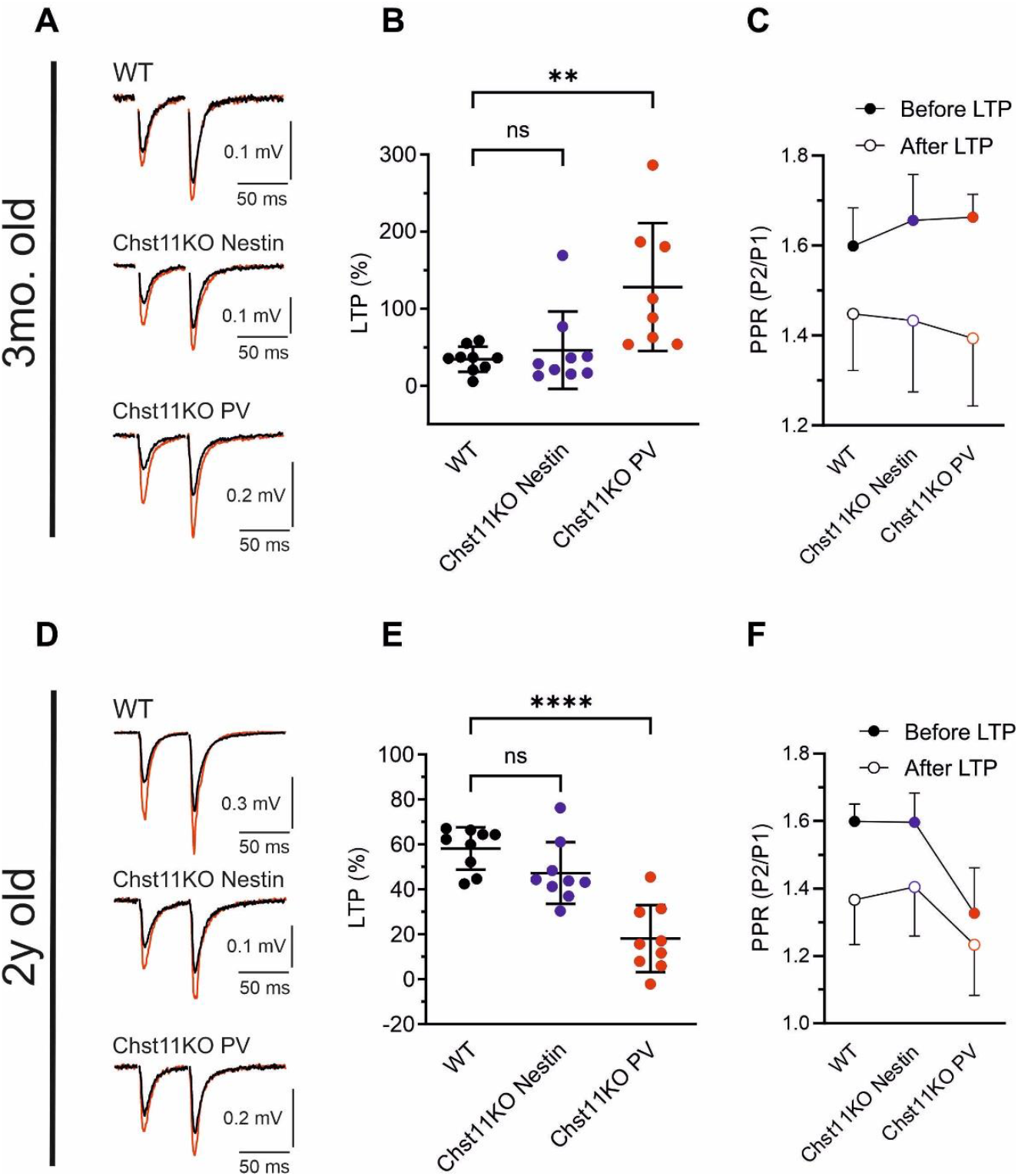
(A) Representative field excitatory postsynaptic potentials (fEPSP) traces from CA1 of three-month-old WT, Chst11KO Nestin, and Chst11KO PV mice. fEPSPs were evoked by paired electrical stimuli delivered every 20 s with a 50 ms inter-stimulus interval. Traces represent averages of 10 consecutive fEPSPs recorded before (black) and after LTP induction (red), shown superimposed. (B) The graph quantifies the magnitude of LTP, expressed as the percentage change in fEPSP amplitude after LTP induction, in 9 WT, 9 Chst11KO Nestin, and 8 Chst11KO PV CA1 slices. (C) The average paired pulse ratio (PPR) calculated from 10 consecutive fEPSPs before and after LTP induction as the ratio of the amplitudes of the second (P2) and first (P1) fEPSPs. Note that LTP induction was accompanied by a decrease in PPR values, suggesting a contribution of a presynaptic mechanism to fEPSP potentiation.(D) Representative recordings of fEPSPs from CA1 of two-year-old WT, Chst11KO Nestin, and Chst11KO PV mice recorded before (black) and after LTP induction (red), shown superimposed. (E) LTP magnitude obtained from 9 WT, 9 Chst11KO Nestin, and 9 Chst11KO PV CA1 slices. (F) The average PPR before and after LTP induction. Data are presented as mean ± SD; one-way ANOVA with Dunnett’s multiple comparisons test.

## Discussion

The present study demonstrates that targeted reduction of chondroitin-4-sulfation (C4S) via conditional deletion of Chst11 produces broad, age-stable enhancements in hippocampal and cortical cognitive functions. By integrating structural, synaptic, electrophysiological, and behavioural analyses across the lifespan, we show that C4S is a crucial regulator of PNN architecture, PV interneuron connectivity, and the expression of age-related cognitive decline. Importantly, this manipulation does not eliminate PNNs, but instead reduces their structural stability in a manner that preserves core scaffolding while enhancing plasticity.

### C4S reduction diminishes PNN structural complexity while retaining somatic organisation

PNNs provide a molecular scaffold that restricts plasticity by consolidating synapses and regulating the excitability of PV interneurons (Miyata & Kitagawa, 2016; Irvine & Kwok, 2018; Rowlands et al., 2018). Here, we show that reducing C4S through Chst11 deletion substantially lowers total aggrecan incorporation into CA1 PNNs and diminishes dendritic PNN coverage, while leaving somatic PNN organisation largely intact. This phenotype contrasts with the more severe loss of PNNs observed with global aggrecan deletion (Rowlands et al., 2018; Ruzicka et al., 2022), quadruple CSPG knockouts (Gottschling et al., 2019; Mueller-Buehl et al., 2022), and link-protein deficiency (Carulli et al., 2010; Romberg et al., 2013). These complete or near-complete ablations disrupt PNN integrity and often induce vulnerability rather than controlled plasticity.

In contrast, Chst11 deletion selectively destabilises the dendritic components of PNNs, consistent with the idea that PNN stability depends not only on core proteins but also on the sulphation pattern of CS-GAG chains. The C6S epitope has been implicated in maintaining a permissive, juvenile-like state by limiting aggrecan retention (Miyata et al., 2012; Miyata & Kitagawa, 2016), while the present findings suggest that C4S plays an opposing role by promoting PNN consolidation. Thus, altering sulphation provides a subtler regulatory strategy that reshapes PNN topology without eliminating the structure.

### Increased synaptic input onto PV interneurons with minimal alteration of basal synaptic transmission

PNN integrity is a major determinant of synapse formation on PV interneurons, which are powerfully regulated by CSPG motif availability, Sema3A binding, and inhibitory structural constraints (Foscarin et al., 2017; Yang et al., 2021). Consistent with this, Chst11 deletion significantly increased excitatory synaptic contacts on PV interneurons across ages, with a genotype-specific enhancement of inhibitory inputs as well, particularly in aged Chst11-PV mice. These results align with earlier reports that PNN disruption promotes formation of new excitatory connections onto PV neurons (Yang et al., 2021) and shifts inhibitory/excitatory ratios following broad CSPG reductions (Gottschling et al., 2019; Mueller-Buehl et al., 2022).

The bidirectional changes observed here—enhanced excitatory connectivity in all *Chst11* knockouts but divergent inhibitory outcomes depending on Cre line—highlight that the role of C4S may differ between developmental and PV-restricted circuits. Mechanistically, reduced aggrecan incorporation may decrease steric hindrance and alter electrostatic properties of synaptic binding surfaces, increasing accessibility for presynaptic terminals.

Electrophysiologically, these synaptic changes produced surprisingly modest functional consequences. *Chst11-PV* ko mice exhibited enhanced LTP at 3 months, consistent with increased excitatory drive onto PV interneurons indirectly facilitating pyramidal plasticity. However, neither knockout exhibited major deviations in basal fEPSPs or LTP at 20 months, contrasting with the substantial fEPSP alterations reported following ChABC treatment (Yang et al., 2015) or Crtl1 knockout (Romberg et al., 2013). The mild phenotype is coherent with our anatomical findings: partial sulphation reduction produces controlled destabilisation rather than collapse of the PNN lattice, preserving the capacity for stable network oscillations.

### Preservation of working memory with aging in the absence of C4S

Working memory decline is an early hallmark of aging and is strongly modulated by PNN maturation, particularly in cortico-hippocampal circuits (Yang et al., 2021). In the spontaneous alternation task, *Chst11-nestin* ko mice maintained high performance through 20 months, whereas WT and *FloxP* animals exhibited age-typical decline beginning at 12 months. The preserved alternation behaviour corroborates the synaptic findings: increased PV synaptic input, without excessive destabilisation, maintains efficient inhibitory control and supports flexible hippocampal encoding.

This phenotype parallels improvements reported with ChABC in the medial prefrontal cortex (Anderson et al., 2020) and with reductions of other restrictive PNN epitopes such as C6S (Yang et al., 2021). The fact that partial C4S depletion produces long-lasting behavioural protection argues strongly that age-related cognitive decline depends on progressive sulphation changes rather than bulk CSPG accumulation alone.

### Enhanced short-term spatial memory across lifespan

Short-term memory performance in the Morris water maze further supports the pro-cognitive role of reduced C4S. *Chst11-nestin* ko mice outperformed controls at 6 months, maintaining superior performance across aging despite increased variability at older ages. The consistency across ages and tasks suggests that C4S reduction enhances hippocampal plasticity in a robust and generalizable manner.

This parallels findings from *Crtl1*ko mice, which exhibit extended memory retention in object recognition and hippocampal-dependent learning (Romberg et al., 2013), as well as from interventions that target C4S itself using Cat-316 antibodies (Yang et al., 2017). The current study adds important nuance by showing that C4S depletion enhances memory without causing global PNN loss or impairing long-term memory consolidation.

### Stable, lifespan-long enhancement of object recognition memory

Object recognition tasks—dependent on perirhinal cortex and supported by hippocampal networks— are particularly sensitive to PNN modifications (Romberg et al., 2013; Yang et al., 2015; 2017; 2021). Both *Chst11-nestin* and *Chst11-PV* ko mice showed enhanced discrimination across all retention delays at 6 and 12 months, consistent with sustained plasticity in perirhinal circuits. Notably, *Chst11-nestin* ko mice retained strong discrimination even at 20 months and 48-h delay, when all other groups declined to chance levels.

This long-lasting enhancement exceeds the duration of memory improvements reported following enzymatic PNN digestion, which typically wane within 24–48 h (Yang et al., 2015; Romberg et al., 2013). As a result, these data position C4S as a critical determinant of memory persistence and highlight *Chst11* modulation as a potential strategy for prolonging cortical memory retention without destabilising network function.

### Improved sociability and preserved social memory in early aging

Social behaviour depends on several PNN-rich regions including hippocampal CA2, where PNNs are essential for social memory encoding (Carstens et al., 2016; Brown et al., 2020). Disruption or abnormal composition of CA2 PNNs leads to sociability deficits (Cope et al., 2021). Here, Chst11 deletion improved sociability and preserved social novelty preference through middle age, whereas WT and *FloxP* animals showed typical age-related decline.

The preservation of sociability across lines suggests that C4S reduction maintains circuit flexibility in CA2 and related limbic regions. However, novelty preference declined in all lines by 20 months. This late-life convergence may reflect aging processes independent of PNN sulphation, such as synaptic loss, microglial activation, or reduced neuromodulatory tone. Nonetheless, the extended window of intact social memory in *Chst11* knockout mice reinforces that C4S contributes to early- and mid-life social cognitive resilience.

### Implications for PNN sulphation as a regulator of aging-related cognitive decline

Multiple studies implicate increases in CSPG C4S and decreases in C6S as hallmarks of age-related synaptic rigidity (Foscarin et al., 2017; Yang et al., 2017; Logsdon et al., 2021). Our findings provide mechanistic evidence that the C4S epitope plays an active role in diminishing plasticity by stabilising aggrecan incorporation and restricting PV interneuron synaptic remodelling. *Chst11* deletion reverses this constraint without eliminating PNNs, resulting in preserved working memory, superior short-term memory, enhanced object memory, and improved sociability.

A critical advantage of sulphation targeting is its subtlety: rather than dissolving PNNs wholesale, modifying their sulphation code allows their structural presence—and their neuroprotective functions—to remain intact. This stands in contrast to enzymatic digestion strategies, which can induce excessive plasticity or destabilise excitatory–inhibitory balance.

### Future directions

The relative contributions of C4S abundance, C6S scarcity, and C4S/C6S ratios to cognitive aging remain unresolved. Given the evidence that altering either epitope can shift plasticity, a combinatorial sulphation code likely dictates PNN function. Further work should characterise region-specific sulphation changes across aging and determine whether targeted modulation can restore plasticity in neurodegenerative contexts where PNN pathology is prominent.

## Acknowledgement

This work was supported by grant from Czech Ministry of Education, Youth and Sports (CZ.02.01.01/00/22_008/0004562) and National Science Foundation 23-05540S.

## Authors contribution

J.S.B., J.S., and J.R. was involved in the experimental design, execution, data analysis and manuscript preparation; J.R. was further involved in the study concept; K.S., T.K., L.G., T.S., N. MV., M. K., and R. T. were involved in execution and data analysis, L.M.U. was involved in execution; P.J., J.W.F. and J.C.F.K. were involved in the study concept, experimental design, data interpretation and manuscript preparation.

## References

Anderson MD, Paylor JW, Scott GA, Greba Q, Winship IR, Howland JG (2020) ChABC infusions into medial prefrontal cortex, but not posterior parietal cortex, improve the performance of rats tested on a novel, challenging delay in the touchscreen TUNL task. Learn Mem 27:222–235.

Bernard C, Prochiantz A (2016) Otx2-PNN Interaction to Regulate Cortical Plasticity. Neural Plast 2016:7931693.

Beurdeley M, Spatazza J, Lee HH, Sugiyama S, Bernard C, Di Nardo AA, Hensch TK, Prochiantz A (2012) Otx2 binding to perineuronal nets persistently regulates plasticity in the mature visual cortex. J Neurosci 32:9429–9437.

Bijata M, Labus J, Guseva D, Stawarski M, Butzlaff M, Dzwonek J, Schneeberg J, Böhm K, Michaluk P, Rusakov DA, Dityatev A, Wilczyński G, Wlodarczyk J, Ponimaskin E (2017) Synaptic Remodeling Depends on Signaling between Serotonin Receptors and the Extracellular Matrix. Cell Rep 19:1767–1782.

Brown LY, Alexander GM, Cushman J, Dudek SM (2020) Hippocampal CA2 Organizes CA1 Slow and Fast γ Oscillations during Novel Social and Object Interaction. eNeuro 7.

Carstens KE, Phillips ML, Pozzo-Miller L, Weinberg RJ, Dudek SM (2016) Perineuronal Nets Suppress Plasticity of Excitatory Synapses on CA2 Pyramidal Neurons. J Neurosci 36:6312–6320.

Carulli D, Pizzorusso T, Kwok JC, Putignano E, Poli A, Forostyak S, Andrews MR, Deepa SS, Glant TT, Fawcett JW (2010) Animals lacking link protein have attenuated perineuronal nets and persistent plasticity. Brain 133:2331–2347.

Carulli D, Broersen R, de Winter F, Muir EM, Mešković M, de Waal M, de Vries S, Boele HJ, Canto CB, De Zeeuw CI, Verhaagen J (2020) Cerebellar plasticity and associative memories are controlled by perineuronal nets. Proc Natl Acad Sci U S A 117:6855–6865.

Cheung ST, Miller MS, Pacoma R, Roland J, Liu J, Schumacher AM, Hsieh-Wilson LC (2017) Discovery of a Small-Molecule Modulator of Glycosaminoglycan Sulfation. ACS Chem Biol 12:3126–3133.

Cope EC, Zych AD, Katchur NJ, Waters RC, Laham BJ, Diethorn EJ, Park CY, Meara WR, Gould E (2021) Atypical perineuronal nets in the CA2 region interfere with social memory in a mouse model of social dysfunction. Mol Psychiatry.

Day P, Alves N, Daniell E, Dasgupta D, Ogborne R, Steeper A, Raza M, Ellis C, Fawcett J, Keynes R, Muir E (2020) Targeting chondroitinase ABC to axons enhances the ability of chondroitinase to promote neurite outgrowth and sprouting. PLoS One 15:e0221851.

Favuzzi E, Marques-Smith A, Deogracias R, Winterflood CM, Sánchez-Aguilera A, Mantoan L, Maeso P, Fernandes C, Ewers H, Rico B (2017) Activity-Dependent Gating of Parvalbumin Interneuron Function by the Perineuronal Net Protein Brevican. Neuron 95:639-655.e610.

Fawcett JW, Kwok JCF (2022) Proteoglycan Sulphation in the Function of the Mature Central Nervous System. Front Integr Neurosci 16:895493.

Fawcett JW, Fyhn M, Jendelova P, Kwok JCF, Ruzicka J, Sorg BA (2022) The extracellular matrix and perineuronal nets in memory. Mol Psychiatry.

Foscarin S, Raha-Chowdhury R, Fawcett JW, Kwok JCF (2017) Brain ageing changes proteoglycan sulfation, rendering perineuronal nets more inhibitory. Aging (Albany NY) 9:1607–1622.

Gogolla N, Caroni P, Lüthi A, Herry C (2009) Perineuronal nets protect fear memories from erasure. Science 325:1258–1261.

Gottschling C, Wegrzyn D, Denecke B, Faissner A (2019) Elimination of the four extracellular matrix molecules tenascin-C, tenascin-R, brevican and neurocan alters the ratio of excitatory and inhibitory synapses. Sci Rep 9:13939.

Happel MF, Niekisch H, Castiblanco Rivera LL, Ohl FW, Deliano M, Frischknecht R (2014) Enhanced cognitive flexibility in reversal learning induced by removal of the extracellular matrix in auditory cortex. Proc Natl Acad Sci U S A 111:2800–2805.

Hou X, Yoshioka N, Tsukano H, Sakai A, Miyata S, Watanabe Y, Yanagawa Y, Sakimura K, Takeuchi K, Kitagawa H, Hensch TK, Shibuki K, Igarashi M, Sugiyama S (2017) Chondroitin Sulfate Is Required for Onset and Offset of Critical Period Plasticity in Visual Cortex. Sci Rep 7:12646.

Huang H, Joffrin AM, Zhao Y, Miller GM, Zhang GC, Oka Y, Hsieh-Wilson LC. Chondroitin 4-O-sulfation regulates hippocampal perineuronal nets and social memory (2023). Proc Natl Acad Sci U S A. 120(24):e2301312120.

Irvine SF, Kwok JCF (2018) Perineuronal Nets in Spinal Motoneurones: Chondroitin Sulphate Proteoglycan around Alpha Motoneurones. Int J Mol Sci 19.

Kwok JCF, Foscarin, S. and Fawcett, J. (2015) Perineuronal nets: a special structure in the central nervous system extracellular matrix. In, pp 23–32. Neuromethods: Springer, New York.

Logsdon AF, Francis KL, Richardson NE, Hu SJ, Faber CL, Phan BA, Nguyen V, Setthavongsack N, Banks WA, Woltjer RL, Keene CD, Latimer CS, Schwartz MW, Scarlett JM, Alonge KM (2021) Decoding perineuronal net glycan sulfation patterns in the Alzheimer’s disease brain. Alzheimers Dement.

Miyata S, Kitagawa H (2016) Chondroitin 6-Sulfation Regulates Perineuronal Net Formation by Controlling the Stability of Aggrecan. Neural Plast 2016:1305801.

Miyata S, Komatsu Y, Yoshimura Y, Taya C, Kitagawa H (2012) Persistent cortical plasticity by upregulation of chondroitin 6-sulfation. Nat Neurosci 15:414–422, S411-412.

Morellini F, Sivukhina E, Stoenica L, Oulianova E, Bukalo O, Jakovcevski I, Dityatev A, Irintchev A, Schachner M (2010) Improved reversal learning and working memory and enhanced reactivity to novelty in mice with enhanced GABAergic innervation in the dentate gyrus. Cereb Cortex 20:2712–2727.

Mueller-Buehl C, Reinhard J, Roll L, Bader V, Winklhofer KF, Faissner A (2022) Brevican, Neurocan, Tenascin-C, and Tenascin-R Act as Important Regulators of the Interplay Between Perineuronal Nets, Synaptic Integrity, Inhibitory Interneurons, and Otx2. Front Cell Dev Biol 10:886527.

Prieur EAK, Jadavji NM. Assessing Spatial Working Memory Using the Spontaneous Alternation Y-maze Test in Aged Male Mice (2019). Bio Protoc. 9(3):e3162.

Romberg C, Yang S, Melani R, Andrews MR, Horner AE, Spillantini MG, Bussey TJ, Fawcett JW, Pizzorusso T, Saksida LM (2013) Depletion of perineuronal nets enhances recognition memory and long-term depression in the perirhinal cortex. J Neurosci 33:7057–7065.

Rowlands D, Lensjø KK, Dinh T, Yang S, Andrews MR, Hafting T, Fyhn M, Fawcett JW, Dick G (2018) Aggrecan Directs Extracellular Matrix-Mediated Neuronal Plasticity. J Neurosci 38:10102–10113.

Ruzicka J, Dalecka M, Safrankova K, Peretti D, Jendelova P, Kwok JCF, Fawcett JW (2022). Perineuronal nets affect memory and learning after synapse withdrawal. Transl Psychiatry. 12(1):480.

Ruzicka J, Kulijewicz-Nawrot M, Rodrigez-Arellano JJ, Jendelova P, Sykova E (2016) Mesenchymal Stem Cells Preserve Working Memory in the 3xTg-AD Mouse Model of Alzheimer’s Disease. Int J Mol Sci 17.

Sahu S, Li R, Loers G, Schachner M. Knockdown of chondroitin-4-sulfotransferase-1, but not of dermatan-4-sulfotransferase-1, accelerates regeneration of zebrafish after spinal cord injury (2019). FASEB J. 33(2):2252–2262.

Saroja SR, Sase A, Kircher SG, Wan J, Berger J, Höger H, Pollak A, Lubec G (2014) Hippocampal proteoglycans brevican and versican are linked to spatial memory of Sprague-Dawley rats in the morris water maze. J Neurochem 130:797–804.

Testa D, Prochiantz A, Di Nardo AA (2019) Perineuronal nets in brain physiology and disease. Semin Cell Dev Biol 89:125–135.

Wingert JC, Sorg BA (2021) Impact of Perineuronal Nets on Electrophysiology of Parvalbumin Interneurons, Principal Neurons, and Brain Oscillations: A Review. Front Synaptic Neurosci 13:673210.

Yang M, Silverman JL, Crawley JN (2011) Automated three-chambered social approach task for mice. Curr Protoc Neurosci Chapter 8:Unit 8.26.

Yang S, Cacquevel M, Saksida LM, Bussey TJ, Schneider BL, Aebischer P, Melani R, Pizzorusso T, Fawcett JW, Spillantini MG (2015) Perineuronal net digestion with chondroitinase restores memory in mice with tau pathology. Exp Neurol 265:48–58.

Yang S, Hilton S, Alves JN, Saksida LM, Bussey T, Matthews RT, Kitagawa H, Spillantini MG, Kwok JCF, Fawcett JW (2017) Antibody recognizing 4-sulfated chondroitin sulfate proteoglycans restores memory in tauopathy-induced neurodegeneration. Neurobiol Aging 59:197–209.

Yang S, Gigout S, Molinaro A, Naito-Matsui Y, Hilton S, Foscarin S, Nieuwenhuis B, Tan CL, Verhaagen J, Pizzorusso T, Saksida LM, Bussey TM, Kitagawa H, Kwok JCF, Fawcett JW (2021) Chondroitin 6-sulphate is required for neuroplasticity and memory in ageing. Mol Psychiatry 26:5658–5668.

Yoshioka N, Miyata S, Tamada A, Watanabe Y, Kawasaki A, Kitagawa H, Takao K, Miyakawa T, Takeuchi K, Igarashi M (2017) Abnormalities in perineuronal nets and behavior in mice lacking CSGalNAcT1, a key enzyme in chondroitin sulfate synthesis. Mol Brain 10:47.

