## Supplementary figures for "Chondroitin 4-sulphate depletion enhances synaptic plasticity and memory in aging"

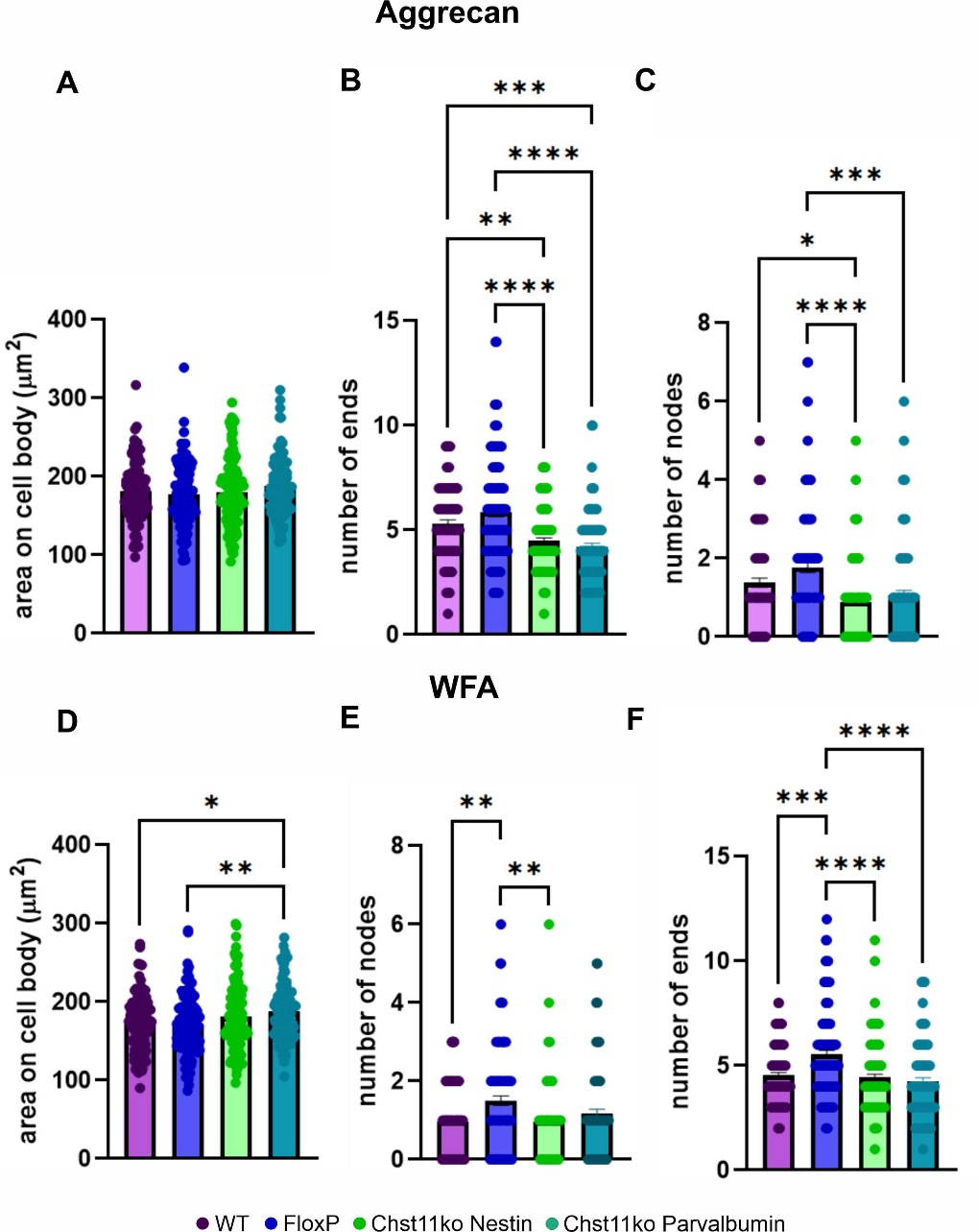

**Figure S1.** Reduction of C4S and immunohistochemistry images showing additional modes of analysis of PNN Structure, including PNN covered cell body area (A, D), number of ends (B, E) and nodes (C, F). The size of PNN covered body area was very similar among the groups. *FloxP* littermate control animals had significantly higher number of ends than *Chst11* knockouts and WT control. Number of nodes was in *Chst11* ko animals significantly reduced especially in aggrecan positive PNNs. *p< 0.05, **p< 0.01, *** p< 0.001, **** p< 0.0001. (One-way ANOVA **A** F (2,343) =2.408 p=0.0915, **B** F (2,343) = 14.02 p< 0.0001, WT vs. *FloxP* q=5.945 p< 0.0001 *FLoxP* vs. *Chst11-nestin* ko q=6.793 p< 0.0001, **C** F (2,343) = 8.908 p= 0.0002, *FloxP* vs. WT q=4.962 p= 0.0015, *FLoxP* vs. *Chst11-nestin* ko q=5.243 p= 0.0007, **D** F (2,337) = 0.2853 p= 0.752, **E** F (2,337) = 15.21 p< 0.0001, WT vs. *Chst11-nestin* ko q=4.494 p= 0.0046, *FLoxP* vs. *Chst11-nestin* ko q=7.746 p< 0.0001, **F** F (2,337) = 15.78 p< 0.0001,WT vs. *FloxP* q=3.34 p=0.049, WT vs. *Chst11-nestin ko* q=4.219 p= 0.0086, *FloxP* vs. *Chst11-nestin* ko q=7.928 p< 0.0001)

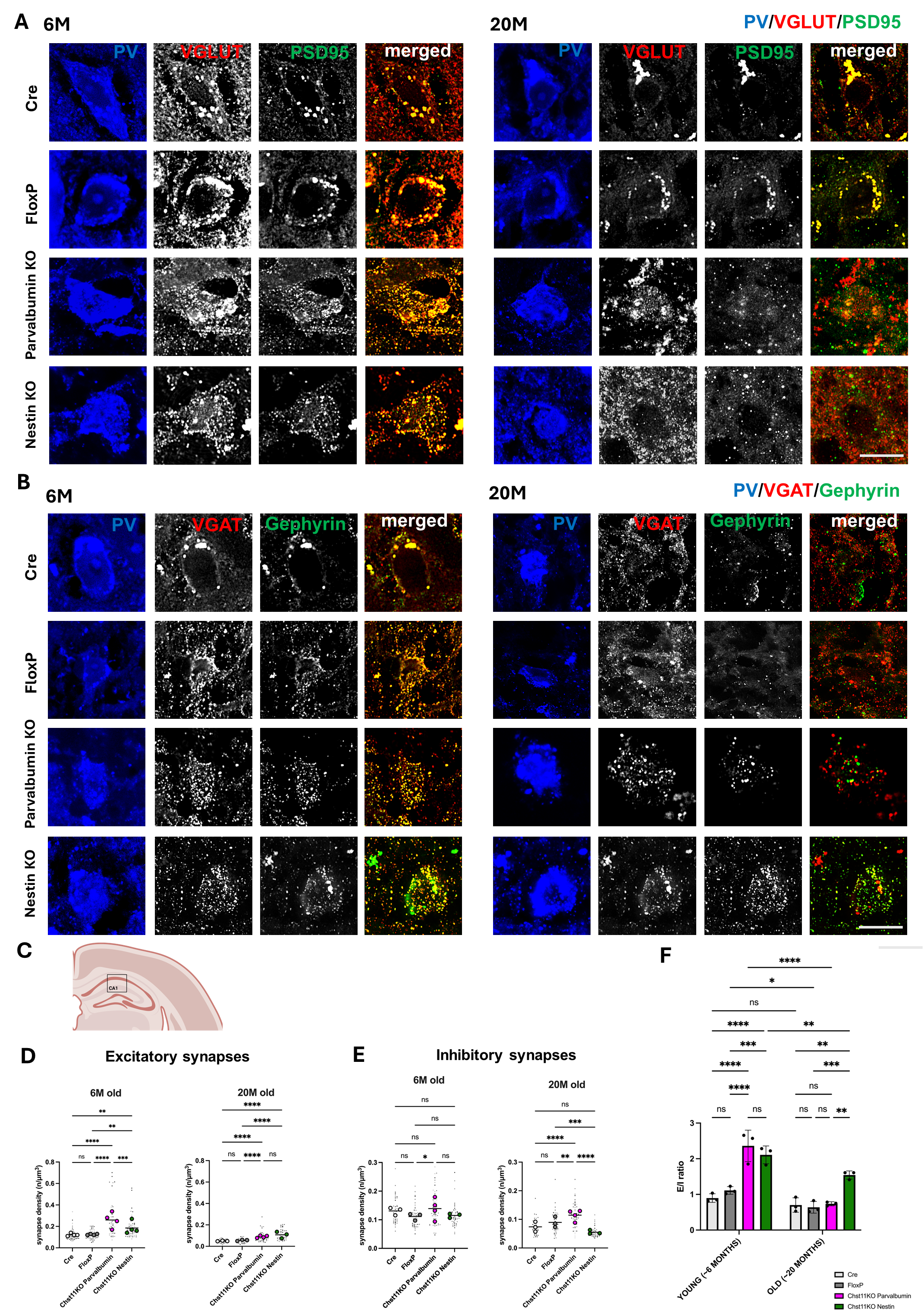

**Figure S2: Synaptic organisation around parvalbumin-positive interneurons in the CA1 region of the hippocampus.** (A, B) Representative confocal images acquired using a Zeiss LSM 880 with Airyscan. The images on the left depict tissue from 6-month-old animals, while the images on the right depict tissue from 20-month-old animals. Scale bar: 10 um.
